# One problem, too many solutions: How costly is honest signalling of need?

**DOI:** 10.1101/240440

**Authors:** Szabolcs Számadó, Dániel Czégel, István Zachar

## Abstract

The “cost of begging” is a prominent prediction of costly signalling theory, suggesting that offspring begging has to be costly in order to be honest. More specifically, it predicts that there is a single cost function for the offspring (depending on e.g. offspring quality) that maintains honesty and it must be proportional to parent’s fitness loss. Here we show another interpretation of the cost. We demonstrate that cost, proportional to the fitness gain of the offspring, also results in honest signalling. Since the loss of the parent does not necessarily coincide with the gain of the offspring, it is provable that any linear combination of the two cost functions (one proportional to parent’s loss, one to offspring’s gain) also leads to honest signalling. Our results, applied for a specific model, support the previous general conclusion that signalling games have different cost functions for different equilibria. Consequently, costly signalling theory cannot predict a unique equilibrium cost in signalling games especially in case of parent-offspring conflicts. As an important consequence, any measured equilibrium cost in real cases has to be compared both to the parent’s fitness loss and to the offspring’s fitness gain in order to provide meaningfully interpretation.

## Background

Parent-offspring communication is a hotly debated topic, continuously in the forefront of behavioural sciences [1–4]. Its appeal stems from the seemingly controversial interests of involved parties. Despite the obvious conflict of interest between parent and offspring [5], offspring frequently solicit food from the parents. In general, this solicitation is found to be honest as more needy offspring begs more intensively [6]. Game theoretical explanations of begging behaviour have gained much attention over the years [7–16]. Most of these game theoretical models predicted costly signalling [7], which became the dominant expectation in past decades.

Nöldeke and Samuelson [17] offered an enlightening account of the cost of honest signalling of need. They have demonstrated that at equilibrium (where honest signalling exists), the signalling cost of the offspring is proportional to the fitness loss of the parent resulting from the transfer of resources. They also showed that the factor of proportionality is solely determined by the degree of relatedness between parent and offspring. Consequently, they claimed that the offspring's condition (and its expected benefit due to the received resource) influences the signalling cost only to the extent that it influences the parent's loss of fitness. Here we extend their model and prove that their solution is not unique and that there is another equilibrium with honest signalling where their claim does not apply, but which can be readily derived from their equations [17]. At this second equilibrium, the cost of signalling is proportional to the expected fitness benefit of the offspring, and (analogously to the other case) the parent's fitness loss affects the signalling cost only to the extent it affects the offspring’s gain. Moreover, any linear combination of these two cost functions provides an equilibrium with honest signalling. Thus, there is an infinite number of distinct equilibria where honest signalling exists.

## Methods

Nöldeke and Samuelson [17] designed their model based on the seminal work of Godfray [7]. They have calculated the fitness functions of the two parties, parent and offspring. The parent is interested in the condition of the offspring to transfer the least amount of resource to maximize its own inclusive fitness (all future offspring included) whereas the offspring is interested in receiving the most amount of resource possible to maximize its own inclusive fitness (all future siblings included). The offspring’s condition is described by a strictly positive continuous variable (*c*). The requirement for signalling stems from the fact that the parent cannot asses this condition directly. The offspring, however, can opt to engage in communication with a (costly) signal (*x*). In the original model of Nöldeke and Samuelson, *x* denoted both the level (intensity) and the cost of the signal [17]. Here, we introduce function *f*(*x*) as the cost of the signal, and reserve *x* to denote only the intensity of the signal (depending on the condition *c*) in order to avoid potential confusion.

The parent has control over *Z* amount of resource that it has to divide between the offspring and itself, where offspring receives part *z* of *Z* and parent retains part *y* = *Z* – *z*. The inclusive fitness functions of offspring and parent (*v* and *u*, respectively, after [17]) are:

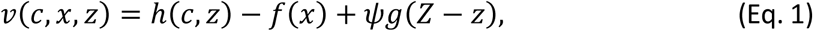

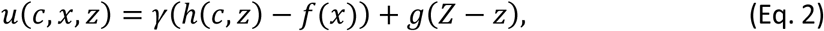

where *h*(*c*, *z*) and *g*(*Z* – *z*) are the direct fitness gains of offspring and parent, respectively, when *z* amount of resource is transferred to offspring. Both *h* and *g* are assumed to be continuously differentiable and increasing functions (accordingly strictly decreasing with *z*). *ψ* is the coefficient of relatedness between current offspring (and any future siblings from the parent); and *γ* is the coefficient of relatedness of the parent to its current (and future) offspring. The offspring strategy is the level of solicitation (*x*) as function of the offspring’s quality (*c*), whereas the parental strategy is the level of shared resource (*z*) as a function of offspring solicitation (*x*).

### Conditions of the honest signalling equilibrium

A stable equilibrium of honest signalling requires three conditions to be met: (*i*) signals must be honest, (*ii*) parents have to respond to signals and (*iii*) the equilibrium must be evolutionarily stable. The latter condition implies that there is a pair of optimal parent and offspring strategies (*z*\*(*x*), *x*\*(*c*)) from which it does not worth departing unilaterally for any of the participants [17]. At an honest equilibrium, parents know the condition of the offspring as their signal of need directly corresponds to their level of need. Thus, the parent’s equilibrium strategy has to maximize the parent’s inclusive fitness *u* for any given *c*, i.e. the following inequality must hold [17]:

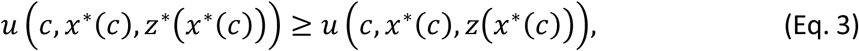

where *x* ^*^ is the equilibrium signal by the offspring, depending on its own quality and *z* ^*^ is the parent’s equilibrium transfer depending on offspring’s signal intensity. Substituting Eq. 2 Into Eq. 3 gives the following condition:

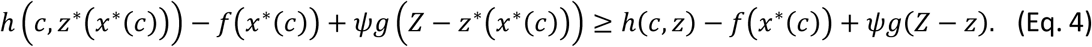

Analogously to parent, offspring’s equilibrium strategy is to maximize its own inclusive fitness *v* given the parental equilibrium strategy *z*^*^(*x*) and the condition of the offspring *c*. Thus, the following condition must hold for any *c* and *x* [17]:

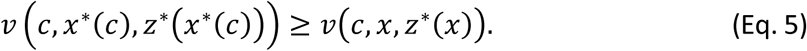

Substituting into Eq. 1 gives the following condition:

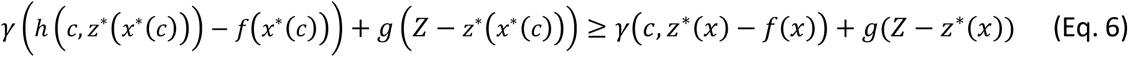

In a signalling equilibrium, the parent’s transfer must satisfy [17]:

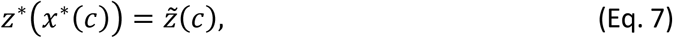

where *x* ^*^ denotes the offspring’s equilibrium signal intensity, *z*^*^ the parent’s equilibrium transfer, and 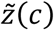 the parent’s optimal transfer.

## Results

The argument of Nöldeke and Samuelson [17] is as follows: the cost of signal at equilibrium has to dispense the conflict of interest between parent and offspring. Accordingly, the two solution functions of *h* and *g* of the optimization problems of parent and offspring have to give the same result (see [18] for more general results). In the absence of signalling cost, at the maximum of the offspring’s inclusive fitness, the following conditions must be met:

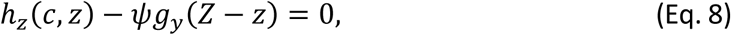

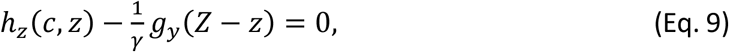

where subscripts denote derivatives with respect to the variable. At the optimum, the derivatives of the two components of the fitness gain must equal:

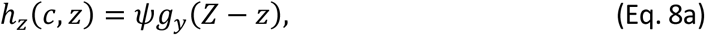

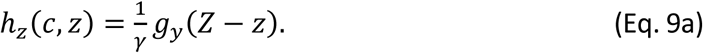

Clearly, the marginal fitness gain of the offspring (in the absence of signal cost) is different from the offspring’s point of view (Eq. 8a) than from the parent’s point of view (Eq. 9a), hence they maximize different functions. Thus, there is a clear conflict of interest between parent and offspring. An illustration of this conflict and the corresponding trade-off can be seen on Figure 1. The shape of these trade-offs is different since the weights of the parental fitness component (*g*) and the offspring fitness component (*h*) are different for the offspring and the parent. The fitness components of the inclusive fitness of the offspring and the parents change alongside the blue and yellow curve respectively with increasing value of *z*. The trade-off implies that one component cannot be increased without the loss of fitness in the other component. Blue and yellow star represents optimal resource allocation and blue and yellow dot indicates the position (fitness) of the offspring and the parent respectively when the resource allocation is optimal for the other party. Clearly the dots do not overlap with the stars, hence the optimal resource allocation of one party is not optimal for the other.

**Figure 1.**
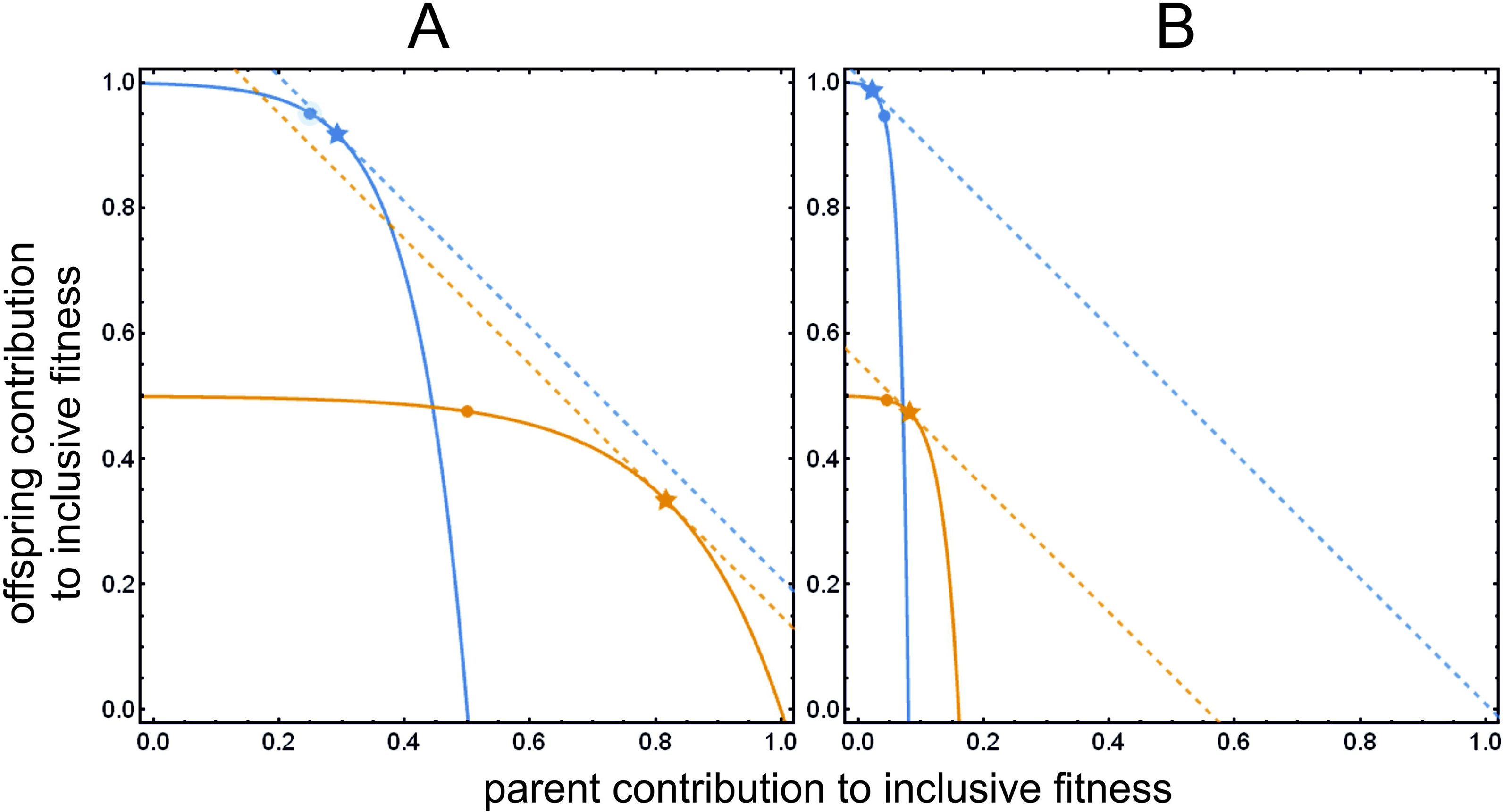
Inclusive fitness functions and optima without signalling cost. **A**: *G* = ½ (it provides a better layout); **B**: *G* = 0.08 (as in [7]). Parent’s (yellow curve, function *u* according to Eq. 2 without signal cost *f*(*x*)) and offspring (blue curve, function *v* according to Eq. 1 without signal cost *f*(*x*) inclusive fitness functions, parameterized by *z*. The *x* coordinate value of parent’s curve is the parent’s own fitness contribution *g*(*c*, *z*), the *y* coordinate value is the fitness contribution the offspring (*γ h*(*c*, *z*)); similarly, the *x* value of the offspring’s curve is the parent’s contribution (*ψ g*(*z*)), the *y* value is the offspring’s own fitness *h*(*c*, *z*). The actual inclusive fitness value is the sum of the appropriate coordinate values, both for parent and offspring. Parameters are *Z* = 2, *γ* = 1/2, *ψ* = ½, *U* = 1, *c* = 3. Yellow and blue stars indicate parent’s and offspring’s optimum. Dashed lines are the calculated derivative tangents that touch optima at 45°, indicating maximum fitness. The optimum *z* value for parent and offspring are not identical: the yellow dot indicates what parent’s fitness at the offspring’s optimum *z*; blue dot is the offspring’s fitness in case of parent’s optimum *z*.

Nöldeke and Samuelson [17] proposed that the cost of signals should resolve this conflict in the honest signalling equilibrium. They [17] proposed the following cost function:

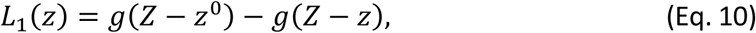

where *z*^o^ is the resource requirement of the offspring in the least needy condition, that is 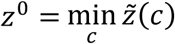 [17]. The cost at equilibrium is:

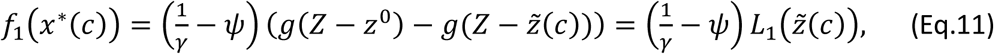

where 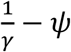 defines the magnitude of the parent-offspring conflict. In equilibrium, 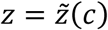. The relationship between *f*(*x*) and *L*(*\**) discussed further in *ESM Appendix 1*.

So far, we have followed the design of Nöldeke and Samuelson [17]. However, starting from the same equations (Eqs. 8 and 9), a different cost function of signalling can also be obtained. Instead of providing the optimality conditions to calculate the offspring’s marginal fitness gain, one can similarly rearrange Eqs. 8 and 9 to calculate the parental marginal fitness gain, from the offspring’s point of view (without signal cost):

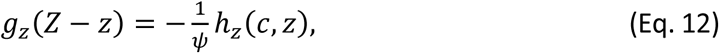

and from the parent’s point of view:

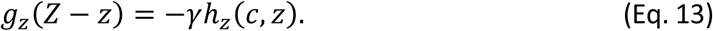

Clearly, in the absence of signal cost, the marginal fitness gain of the parent (as a function of resource allocation) is different from the offspring’s point of view (Eq. 11) than from the parent’s point of view (Eq. 12). This still implies the conflict of interest. Following the same logic as above, at the honest signalling equilibrium, these equations have to provide the same results. That is, the parent’s optimum has to be the same, viewed either from the offspring’s or from the parent’s aspect. Thus, just as before, the difference between the right-hand sides of Eqs. 11 and 12 gives the cost that has to be subtracted from the offspring fitness so that the two equations result in the same optimum. The cost function we propose is:

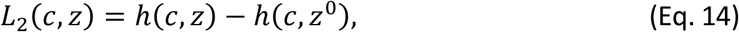

and the cost at equilibrium is:

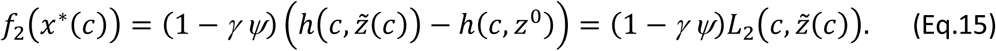

The existence of the signalling equilibrium can be proved as before (see *ESM Appendix 2*).

So far, we have proved that there are two honest signalling equilibria corresponding to two different cost functions. Since each of these cost functions can remove the conflict of interest between parent and offspring, it follows that any linear combination of the functions is also a solution to the optimization problem. Thus, the general cost function of the optimum strategies is as follows:

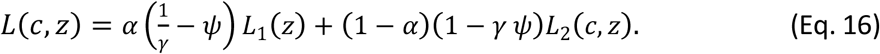

The cost at equilibrium is:

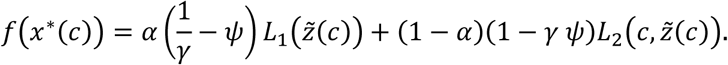

Finally, we provide a numerical example using Godfray's [7] equations. Godfray used the following equations for the offspring’s and parent’s fitness contributions, respectively [7]:

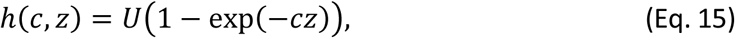

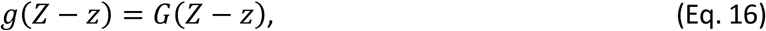

where *U* and *G* are constants. From now on, we use the values provided by Godfray [7]: *U* = 1, *G* = 0.08. Figure 1 shows the actual inclusive fitness values for parent and offspring (*u* of Eq.2 and *v* of Eq. 1, respectively) when there is no cost of signalling, as functions of the quality of the offspring *c* and parental resource allocation *z*. Figure 2 also shows the equilibrium transfer function for parent and for offspring (red curve), which corresponds to the optimal resource allocation for the offspring and the parent, respectively (as a function of *c*). Figure 2 clearly demonstrates that the optima are at different *z* values for the two parties.

**Figure 2.**
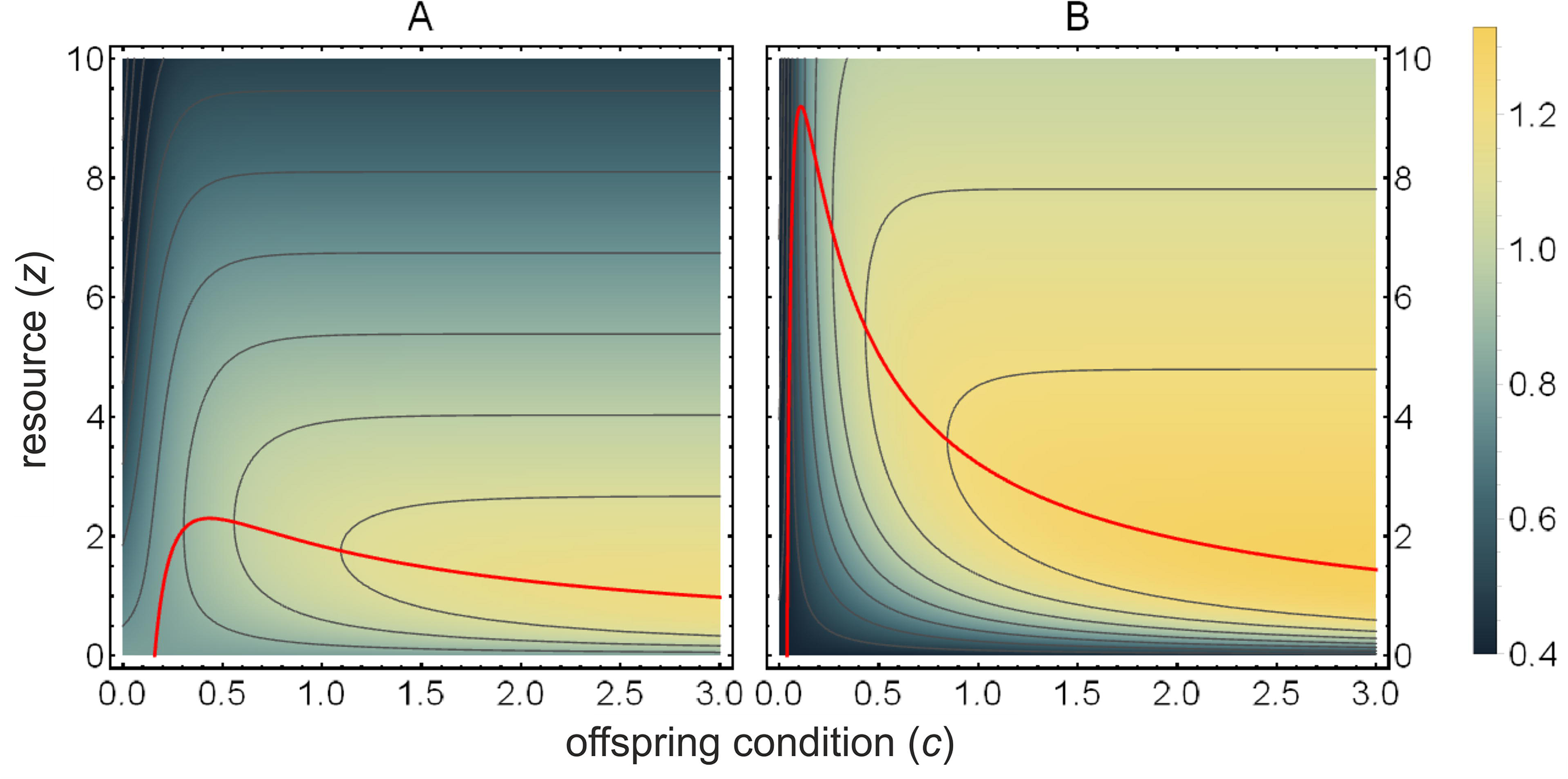
**Inclusive fitness** depending on offspring condition *c* and parental investment *z*. **A**: parental fitness. **B**: offspring fitness. Red line connects the equilibrium *z* values where 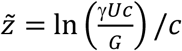 holds. Parameters are *Z* = 10, *G* = 0.08, *U* = 1, *γ* = ½, *ψ* = ½.

Substituting Godfray’s equation (Eqs. 15, 16) into the cost function defined by *L_1_* (Eq. 10) results in:

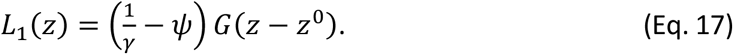

Substituting the same equations into the cost function defined by *L*_2_ (Eq. 13), results in:

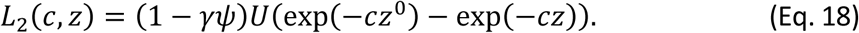

Figure 3 show the same trade-off as Figure 1 just with cost function added to the offspring inclusive fitness (Eqs 1,2). Figure 3A show the new cost function, Figure 3C shows the cost function proposed by Nöldeke and Samuelson [17], while Figure 3B shows a linear combination of the two functions (*α*=0.5). The dots overlap with the stars, hence these cost functions indeed remove the conflict of interest between parent and offspring. We provide an interactive version in the *Electronic Supplementary Material* (*ESM_interactive_figure.nb* and *ESM_interactive_figure.cdf*) that can be used to explore parameter ranges with or without signal cost as well as different linear combinations of these cost functions (*ESM_interactive_figure_video.mp4* shows examples).

**Figure 3.**
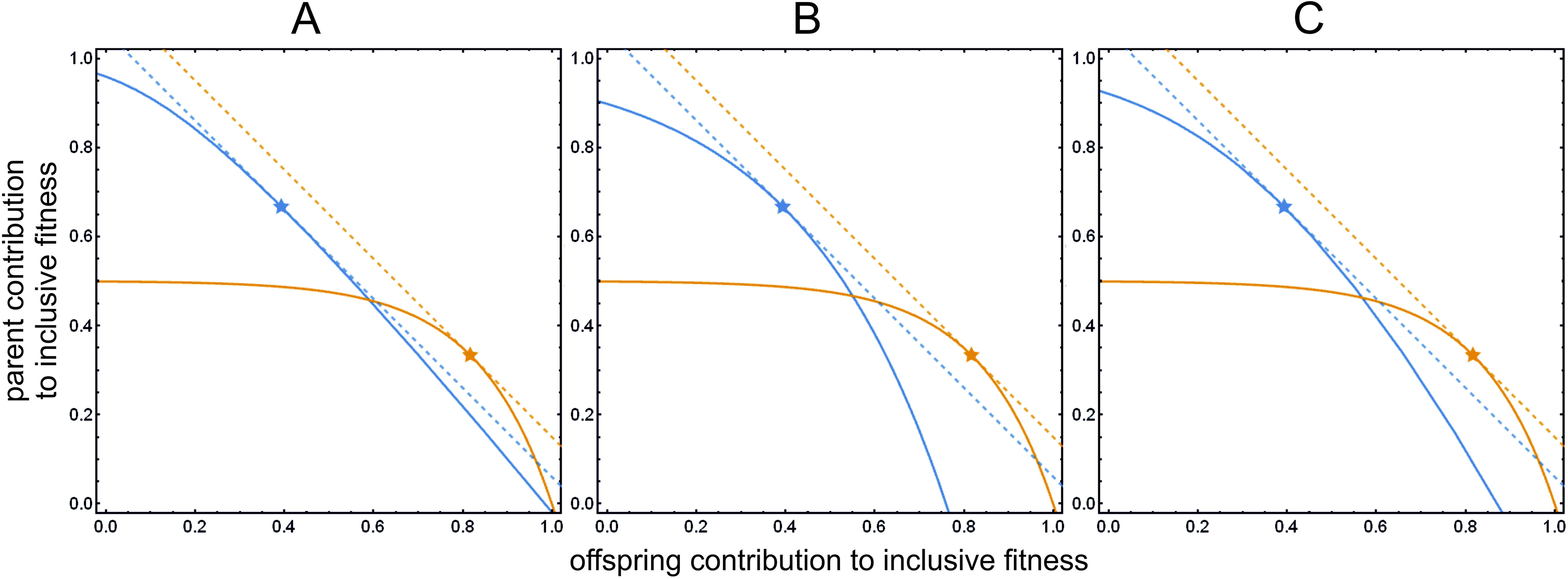
Inclusive fitness functions and optima with signalling cost. Parent’s (yellow curve, function *u* according to Eq. 2) and offspring (blue curve, function *v* according to Eq. 1) inclusive fitness functions, parameterized by *z*. The cost function of Eq. 14 is used with various *α* and *β* values. The *x* coordinate value of parent’s curve is the parent’s own fitness contribution *g*(*c*, *z*), the *y* coordinate value is the fitness contribution of all future offspring (*γ h*(*c*, *z*)); similarly, the *x* value of the offspring’s curve is the parent’s contribution (minus cost) (*ψ g*(*z*)– *f*(*α*, *c*, *z*)), the *y* value is the offspring’s own fitness *h*(*c*, *z*). The actual inclusive fitness value is the sum of the appropriate coordinate values, both for parent and offspring. Parameters are *Z* = 2, *γ* = ½, *ψ* = ½, *U* = 1, *G* = ½, *c* = 3. Yellow and blue stars indicate parent’s and offspring’s optimum. Dashed lines are the calculated derivative tangents that touch optima at 45°, indicating maximum fitness. The optimum *z* value for parent and offspring are always identical, regardless of *α* and *β* values. **A**: Cost function *f*_1_ of Nöldeke and Samuelson [17] (*α* = 1). **B**: Cost function *f*_2_ introduced in this paper (*α* = 0). **C**: Linear combination of the above two cost functions *f*_1_ and *f*_2_(*α* = ½).

Figure 4 shows the actual values for the different cost functions *L*_1_ (Figure 4B), *L*_2_ (Figure 4D) and their linear combination (Figure 4C), when using Godfray’s equation (Eqs. 15, 16). Red, yellow and green curves show the signal cost along the equilibrium path (*f*_1_(*x\**(*c*)) and *f*_2_(*x\**(*c*))). This cost can be calculated by substituting *z* with the amount of optimal parental investment 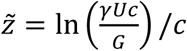 into Eqs. 17 and 18. Figure 4A shows how these equilibrium costs compare to each other as a function of offspring quality *c*. Note, while the absolute value of the equilibrium signal cost is different for each cost function but the partial derivative with respect of *z* is the same along the equilibrium path (see Figure 4F, G and H). Figure 4E illustrates this effect.

**Figure 4.**
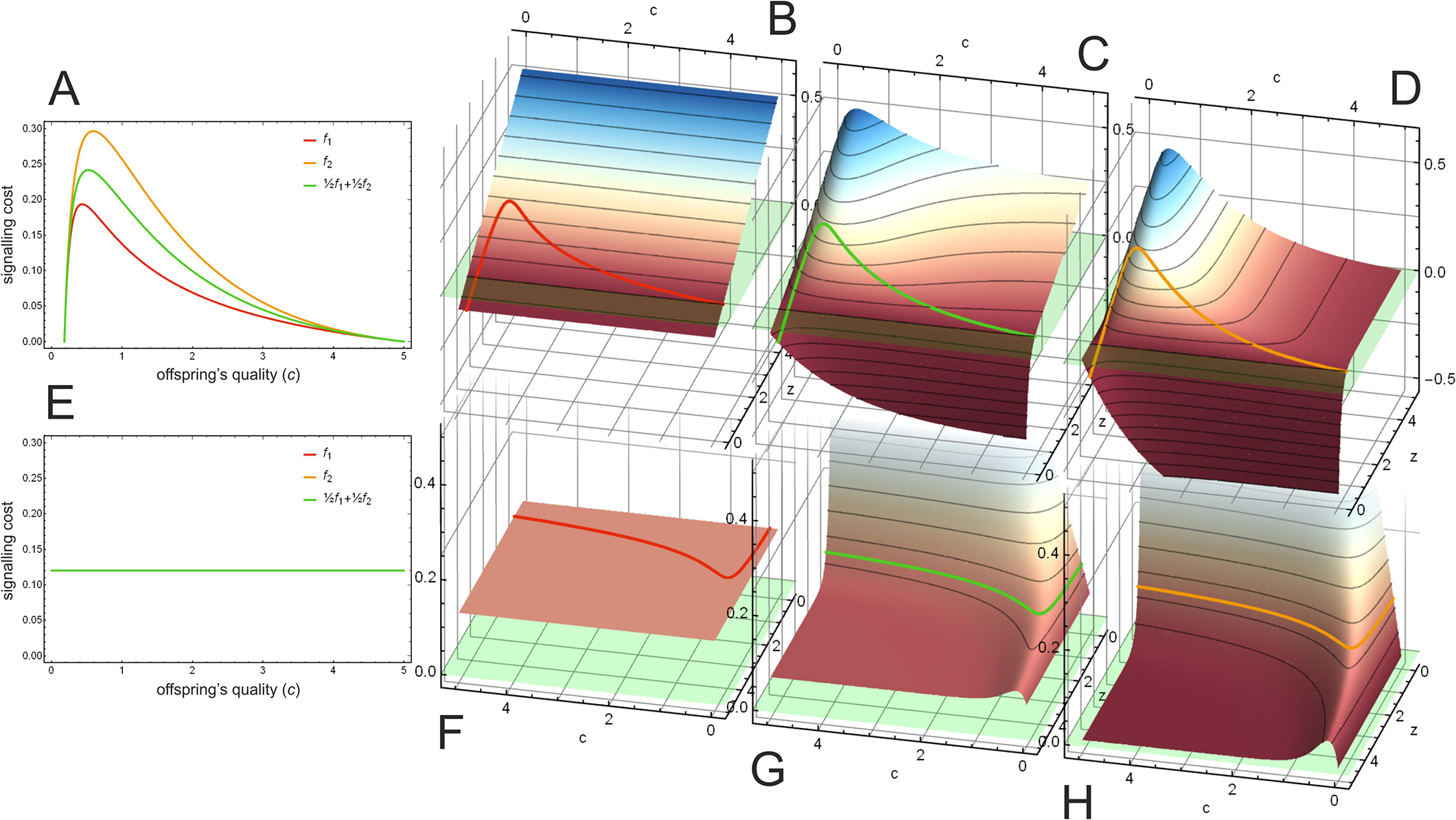
**Signalling cost functions**, depending on offspring condition *c* and parental transfer *z*. **A**: equilibrium signal cost (*f*(*x\**(*c*))) for the different cost functions. **B**: Signal cost function *L*_1_ (Eq. 10). **C**: Linear combination of ½ *L*_1_ + ½ *L*_2_ (Eq. 14) **D**: signal cost function *L*_2_ introduced in this paper (Eq. 13). Red, green and orange curves describe the signal cost (*f*(*x\**(*c*))) along the equilibrium path (which describes the equilibrium transfer function 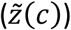 for parent as function of *c*), these curves are shown in panel A. **E**: the partial derivative of the signal cost functions with respect to *z* along the equilibrium path as a function of *c*. **F**: the partial derivative of signal cost function *L*_1_ (Eq. 10) with respect to *z*. **G**: the partial derivative of the linear combination of ½ *L*_1_+½ *L*_2_ (Eq. 14) **H**: the partial derivative of signal cost function *L*_2_ introduced in this paper (Eq. 13) with respect to *z*. Red, green and orange curves describe the partial derivative of the respective signal cost functions along the equilibrium path with respect to *z*, these curves are shown in panel **E**. Parameters are *Z* = 10, *G* = 0.08, *U* = 1, *γ* = ½, *ψ* = ½.

## Discussion

According to Nöldeke and Samuelson [17] (and Eq. 13), based on Godfray’s original differential benefit model [7], the cost of honest signalling should be proportional to parent’s fitness loss. But is it the only solution that yields honest signalling in equilibrium? While the existence of infinite costly equilibria is known in general [18], no other equilibrium has been calculated yet in terms of Godfray’s model. Here, we show that a second extremum exists when the cost is proportional to offspring's fitness gain, which also yields a signalling equilibrium with costly signals. Furthermore, we have demonstrated, that any linear combination of the two extreme cost functions is an equilibrium itself, which effectively proves that an infinite number of honest, evolutionarily stable costly signalling equilibria exist for Godfray’s model. While we specifically derived the second extremum cost function for Godfray’s model, our results have important theoretical and empirical implications.

There are six major theoretical outcomes concerning the signalling of need, which apply generally: (*i*) the population is not in equilibrium [16]; (*ii*) there is, on average, a shared interest between parent and offspring, hence partially honest pooling equilibria can exist with cost-free signals [11, 13]; (*iii*) there is an honest signalling equilibrium in a differential benefit model [7], where the cost of signalling is proportional to the parent's fitness loss [17]; (*iv*) as we have shown, there exists an honest signalling equilibrium in a differential benefit model [7], where the cost of signalling is proportional to the offspring's expected fitness gain; (*v*) there is an infinite number of honest signalling equilibria where the cost of signalling is proportional to the linear combination of the cost functions of the previous two cases including equilibria where the cost of signalling is smaller – even negative for some signallers – than in any other equilibria; and finally (*vi*) it is possible, that a differential cost model offers a better fit for parent-offspring communication (marginally mentioned in [19]. This could open up possibilities for other cost-free [20–22] or even negative cost equilibria [21].

Another important implication of our results and the above considerations is that it is not possible to decide in case of a real population (based on game theoretical models alone) which one of the infinite numbers of costly honest equilibria will be achieved (provided that an honest separating equilibrium exists). In order to answer questions of which evolutionary trajectory will be played out (or have been taken), a more dynamic approach is needed [10]. Godfray and Johnstone [10] calculate the fitness advantage of the signalling equilibrium to the non-signalling equilibrium using the cost function of Nöldeke and Samuelson [17]. Our results could significantly change the outcome of these types of calculations, affecting seriously the evolutionary consequences. This is left for future work.

Since the publication of Godfray’s [7] influential model, a lot of empirical research has been carried out to measure the “cost of begging”. It was realized very early that the metabolic cost of begging is not unreasonably high [23–25], and thus it probably does not fit the predictions of costly signalling theory. Attempts to try to measure the cost of increased begging on growth provided mixed results [26–28]. However, several types of other costs were proposed, like predation risk [29–31], immunological [32–34] or oxidative costs [35]; for a review, see [36]. We must emphasize, that measuring any cost *in absolute value* is not enough [22, 37]: the measured costs have to be *compared* to something, i.e. only relative measures are informative. One of the reasons why the current empirical results are inconclusive is that we don’t have any information about how these costs relate to the benefits of the parties, though see Moreno-Rueda and Redondo [34] for an exception. The results of Nöldeke and Samuelson [17] and the results presented here, alongside with other theoretical results [21, 22], give us a guide how one can meaningfully compare the costs of the different parties involved. It follows, that researchers of the field have to take into account (that is: measure) both the potential fitness loss of the parent *and* the potential fitness gain of the offspring in parent-offspring communication when testing the predictions of costly begging models.

## Author’s contributions

Sz.Sz. conceived the idea; Sz.Sz., D.C. and I.Z. analysed the model; I.Z. analysed the numerical examples and created the figures; all three authors contributed to the writing of the paper.

## Acknowledgements

We would like to thank József Garay for helpful comments.

## Electronic Supplementary Material

ESM Appendix 1–2 ESM Interactive Figure Mathematica notebook ESM Interactive Figure Mathematica interactive computable document format ESM Interactive Figure Video

